# dbGSRV: a manually curated database of genetic susceptibility to respiratory virus

**DOI:** 10.1101/2021.12.26.474200

**Authors:** Ping Li, Yan Zhang, Wenlong Shen, Shu Shi, Zhihu Zhao

**Affiliations:** Beijing Institute of Biotechnology, Beijing, China

## Abstract

Human genetics has been proposed to play an essential role in inter-individual differences in respiratory virus infection occurrence and outcomes. To systematically understand human genetic contributions to respiratory virus infection, we developed the database dbGSRV, a manually curated database that integrated the host genetic susceptibility and severity studies of respiratory viruses scattered over literatures in PubMed. At present, dbGSRV contains 1932 records of genetic association studies relating 1010 unique variants and seven respiratory viruses, manually curated from 168 published articles. Users can access the records by quick searching, batch searching, advanced searching and browsing. Reference information, infection status, population information, mutation information and disease relationship are provided for each record, as well as hyper links to public databases in convenient of users accessing more information. In addition, a visual overview of the topological network relationship between respiratory viruses and associated genes is provided. Therefore, dbGSRV offers a promising avenue to facilitate researchers to dissect human factors in respiratory virus infection, define novel drug targets, conduct risk stratification of population and develop personalized medicine approaches.

Database URL: http://www.ehbio.com/dbGSRV/front/

## Introduction

Respiratory viruses are viruses that enter from respiratory tract and proliferate in respiratory mucosal epithelial cells, causing local infection in respiratory tract or lesions in other organs (1). Common human respiratory viruses include respiratory syncytial virus (RSV), rhinovirus, influenza virus, parainfluenza virus, human metpneumonia virus, coronavirus, adenovirus and so on (2). Respiratory virus infection is one of the leading causes of human mortality and morbidity, which confers constant public health treats and results in significant economic losses (3–5). Of note, as of 25th August, 2021, severe acute respiratory syndrome coronavirus 2 (SARS-CoV-2), a novel coronavirus emerged in 2019 (6–8), has spread all over the world and caused over 213 million infections and 4.4 million deaths (https://covid19.who.int/).

Human responses to respiratory virus infection differ from uninfected, asymptomatic, mild, moderate, severe to fatal outcome. The wide variations in susceptibility and severity are not only attributed to the different transmissibility and virulence of different virus strains, but also attributed to host factors like age, sex, premature birth, pregnancy, obesity and comorbidity (9–11). Among host factors, host genetic background attracts more and more attention in these years (12, 13). Adoption, twin and heritability studies provided the first line of evidence (14–16), followed by candidate-gene study, genome-wide association study (GWAS), whole exome sequencing (WES) and whole genome sequencing (WGS) in recent years (17), revealing that human genetic variants play an important role in susceptibility and severity to infection by altering the expression or function of genes, especially those in genes involved in viral life cycle, host inflammatory and immune response (18–21). These genetic association studies may help dissect the underling mechanisms of viral pathogenesis and host antiviral defense and may contribute to future clinical risk prediction models, allowing for the stratification of individuals according to risk so that those at high risk would be prioritized for immunization (22).

Though great progress has been made in this area, there is a lack of database systematically collecting, formatting, annotating, storing and displaying studies of human susceptibility and severity in respiratory virus. Searching and reading related papers scattered in PubMed are time-consuming, hindering convenient access to useful information.

Therefore, we present the first database of Genetic Susceptibility to Respiratory Virus (dbGSRV), which integrates published genetic studies relating susceptibility and severity in respiratory virus infection. It contains 1932 records of genetic association studies relating 1010 unique variants and seven respiratory viruses, manually curated from 168 published articles. Comprehensive information about reference, infection, samples, mutations and their relationships are available at http://www.ehbio.com/dbGSRV/front/. We anticipate that this resource will a useful tool for researchers to query and retrieve genetic association studies of respiratory viruses.

## Materials and Methods

### Publications collection

We searched for literatures that describe genetic associations with susceptibility or severity of respiratory virus infections in PubMed, using keywords of ‘variant’, ‘polymorphism’, ‘susceptibility’ combined with names of specific respiratory virus like ‘adenovirus’, ‘bocavirus’, ‘influenza’, ‘measles’, ‘MERS’, ‘metapneumovirus’, ‘mumps’, ‘parainfluenza virus’, ‘respiratory syncytial virus’, ‘rhinovirus’, ‘rubella’, ‘SARS’ and ‘SARS-CoV-2’. The searching results were manually examined to only leave English publications that study associations between human single nucleotide variants (SNVs), multiple nucleotide variants (MNVs) or indels with susceptibility or severity of explicit respiratory virus in case-control researches. As for these related publications, we collected publication information like paper title, first and corresponding author, year and journal published and PubMed Unique Identifier (PMID).

### Data extraction, standardization and annotation

We defined one record of genetic association study based on the virus type, case-control sample and variant. As for each record, respiratory virus type information was extracted from the full text of the paper. Virus subtype was also extracted if specified. The number, country, ethnicity and clinical severity information of samples were collected. Based on the ethnicity, we determined the superpopulation of 1000 Genomics that the sample population belonged to. If the sample belonged to multiple superpopulation, marked as ‘Mixed’. No matter whether the variant was associated with virus susceptibility or not, all the studied variants mentioned in the main text were included, to provide a more comprehensive and unbiased scope of genetic association studies.

For sake of uniformity, the name, reference allele and alternate allele of each variant were based on the dbSNP database. Many early publications did not offer dbSNP rs ID of the variant. We manually annotated the rs ID of these variants by genomic mapping. The original names of these variants in the publication were also included in the database as old name. The genomic position of the variants was annotated based on hg38 human genome. Annotation of variants relative to genes were based the following order: exon, 5’ UTR, 3’ UTR, intron, promoter (within 2kb upstream), upstream (2-5kb upstream), downstream (within 2kb downstream).

The alternate allele frequencies of cases and controls, statistic method, odds ratio (OR), 95% confidence interval (CI) and p value for the allele association were extracted from the full-text or supplementary materials. If only genotype frequency was given in the paper, then alternate allele frequency was calculated manually. As for p value, ‘> 0.05%’ was marked if the paper did not give a specific value but claimed that there was no statistically significant difference or association. The allele, genotype, and haplotype association results were each classified into one of the following four categories: ‘severity’, ‘susceptibility’, ‘no association’ and ‘NA’. If at least of one of the allele, genotype or haplotype association result reported ‘severity’ or ‘susceptibility’, then the overall association status was determined as ‘severity’ or ‘susceptibility’, otherwise the overall association status was determined as ‘no association’. Additional noteworthy information about sample, variant and disease association was included in notes.

### Analysis of associated genes

Gene Ontology (GO) and pathway analysis of associated genes were conducted using Database for Annotation, Visualization and Integrated Discovery (DAVID, https://david.ncifcrf.gov/).

## Results

### Web interface

The dbGSRV database comprises six pages, including Home page, Browse page, Batch Search page, Advanced Search page, Network page and Help page.

On the Home page, users can find a brief introduction, update log of dbGSRV and a quick search box (Fig 1A). The quick search box allows users to search genetic association records based on virus name, variant rs ID, gene name or genomic region. The Batch Search page allow users to search multiple viruses, variants, genes, or genomic loci either by entering keywords in the text box or by uploading a txt file (Fig 1B). On Advanced Search page, users can search by logical combination of more keywords (Fig 1C). The Browse page permits users to browse all records by virus, annotation or study type (Fig 1D).

**Fig 1.**
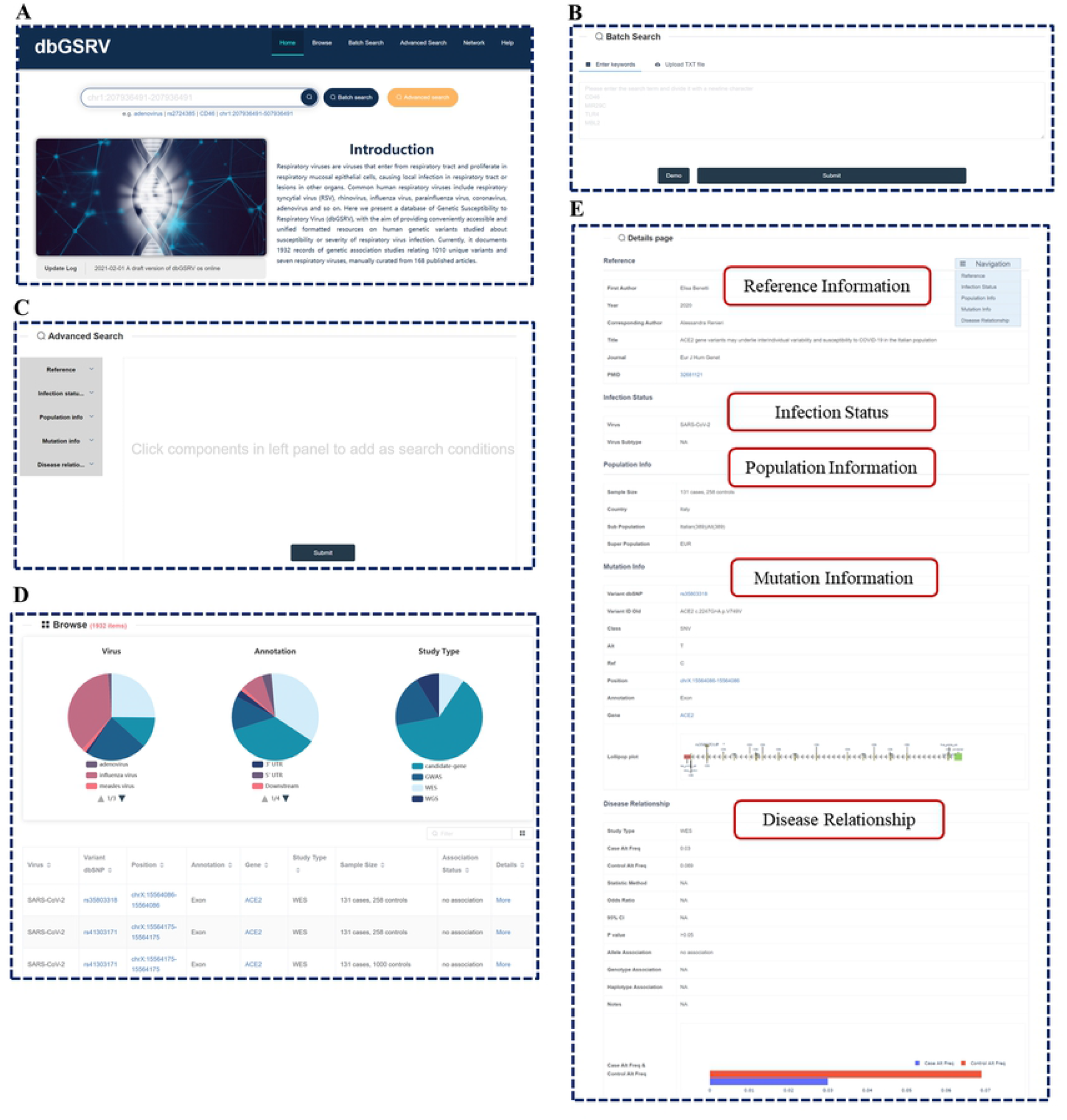
Screen shot of dbGSRV contents. (**A**) Home page. (**B**) Batch Search page. (**C**) Advanced Search page. (**D**) Browse page. (**E**) Detailed information about the record.

The search results are presented as pie charts and a table (Fig 1D). The pie charts display the number and proportion of each subgroup as for virus type, variant position relative to genes and study type respectively, while the table contains the basic information of each record, including virus type, variant information (rs ID in dbSNP database, position in hg38 genome and relative position to genes), study type, sample size and association status. Clicking the subgroup in the pie charts will get the results of the subgroup, and clicking the same subgroup one more time will return back.

In addition, the table provides several features. First, users can further filter the results in the table by typing terms in the ‘filter’ box at the top-right of the table. Second, Clicking the icon on the right of ‘filter’ box, users can change the columns displayed in the table. The following three columns can also be added: ID (unique ID of each record), Year (the year that the paper is published) and PMID (the PMID of the paper in PubMed database). Third, each column could be ranked in ascending or descending by clicking the triangle on the right side of the column header. Fourth, for each record, clicking the Variant dbSNP, Position, Gene and PMID column will take users to the corresponding page in the dbSNP, UCSC, GeneCards and PubMed database respectively. Clicking ‘More’ will jump to the Details page of the record in the database, which consists of more detail information about the reference, infection, population, mutation and disease relationship (Fig 1E).

The Network page provides a visual overview of the topological relationship between respiratory viruses and associated genes (Fig 2). Nodes represent respiratory viruses and genes. Respiratory virus node and gene node are linked by an edge if at least one variant on the gene are reported to be associated with the susceptibility or severity of the virus. As default, the network shows all the respiratory viruses in the database. Users can select a set of specific viruses and submit to generate a new network for these viruses. The attributes of nodes (such as size, shape, background color, label font size and label color) and the overall layout of the network can be edited. The picture can be exported in SVG format for publication usages, as well as the data used to generate the network, which can be downloaded in an Excel file.

**Fig 2.**
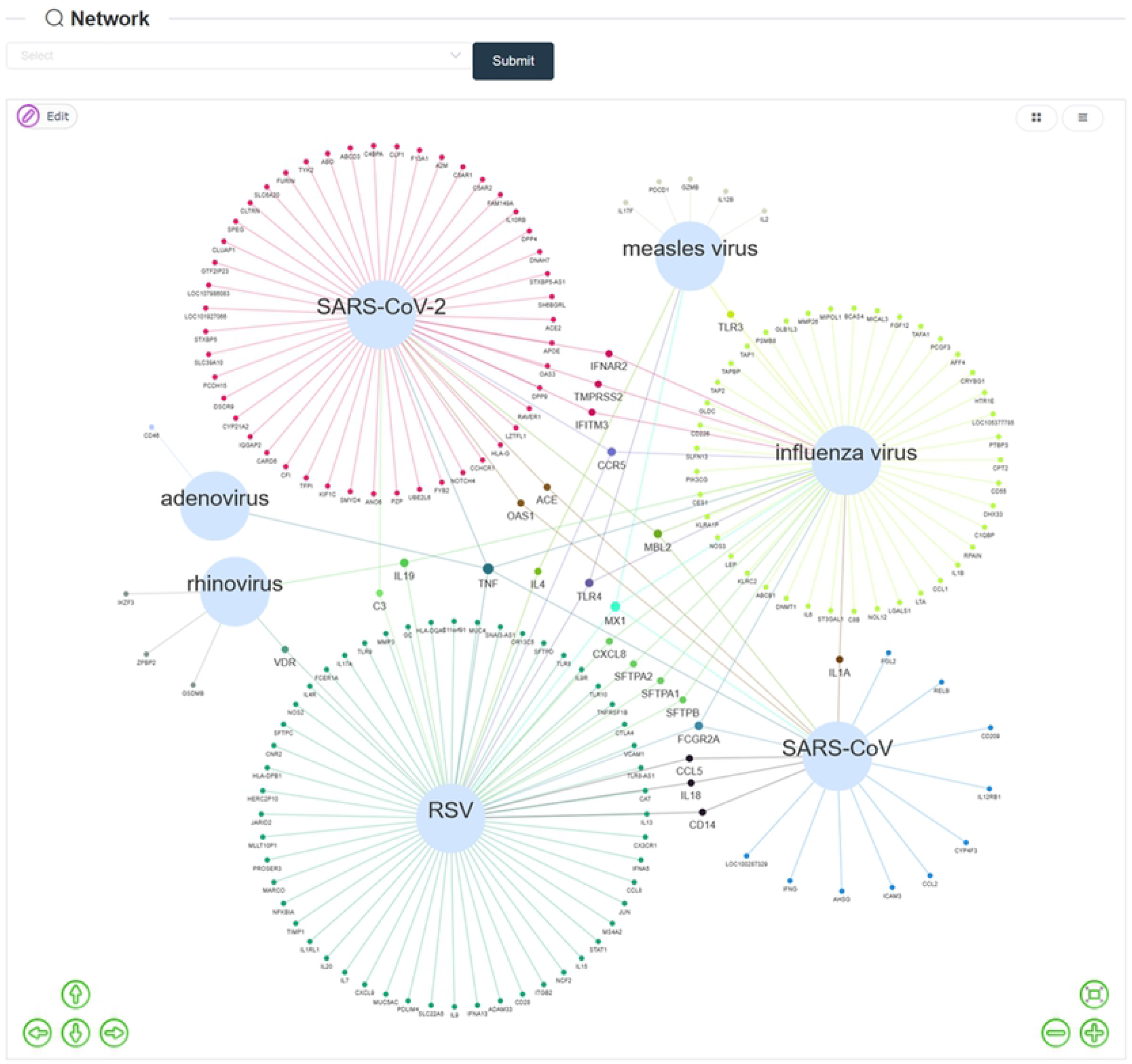
Network page provides a visual overview of the topological relationship between respiratory viruses and associated genes. Nodes represent respiratory viruses and genes. Respiratory virus node and gene node are linked by an edge if at least one variant on the gene are reported to be associated with the susceptibility or severity of the virus. Genes associated with more than one virus are highlighted by larger label size.

At last, dbGSRV provides a detailed tutorial for usage of the database in the Help page.

### Database statistics

For a more comprehensive and unbiased understanding of the genetic association studies with respiratory viruses, the database not only includes positive results of association, but also includes negative results reported in the main text of the paper.

In total, dbGSRV contains 1932 records of genetic association studies relating 1010 unique variants and seven respiratory viruses, manually curated from 168 published articles. The seven respiratory viruses are adenovirus, influenza virus, measle virus, rhinovirus, RSV, SARS-CoV and SARS-CoV-2, of which influenza virus, SARS-CoV-2, RSV and SARS-CoV have the most records, with 718, 486, 433 and 221 records respectively (Fig 3A). Besides, a majority of the records are related with variants residing in the intron (35.7%) and exon (34.3%) of genes (Fig 3B). As for study strategy, most of the records are curated from candidate-gene study (62.4%) (Fig 3C). It is worth noting that 610 records report positive genetic associations between 249 unique variants of 159 genes with respiratory virus infection, mostly based on allele frequency (Fig 3D). Among the positive associations, 149 records are related to susceptibility to infection while 461 records are related to severity.

**Fig 3.**
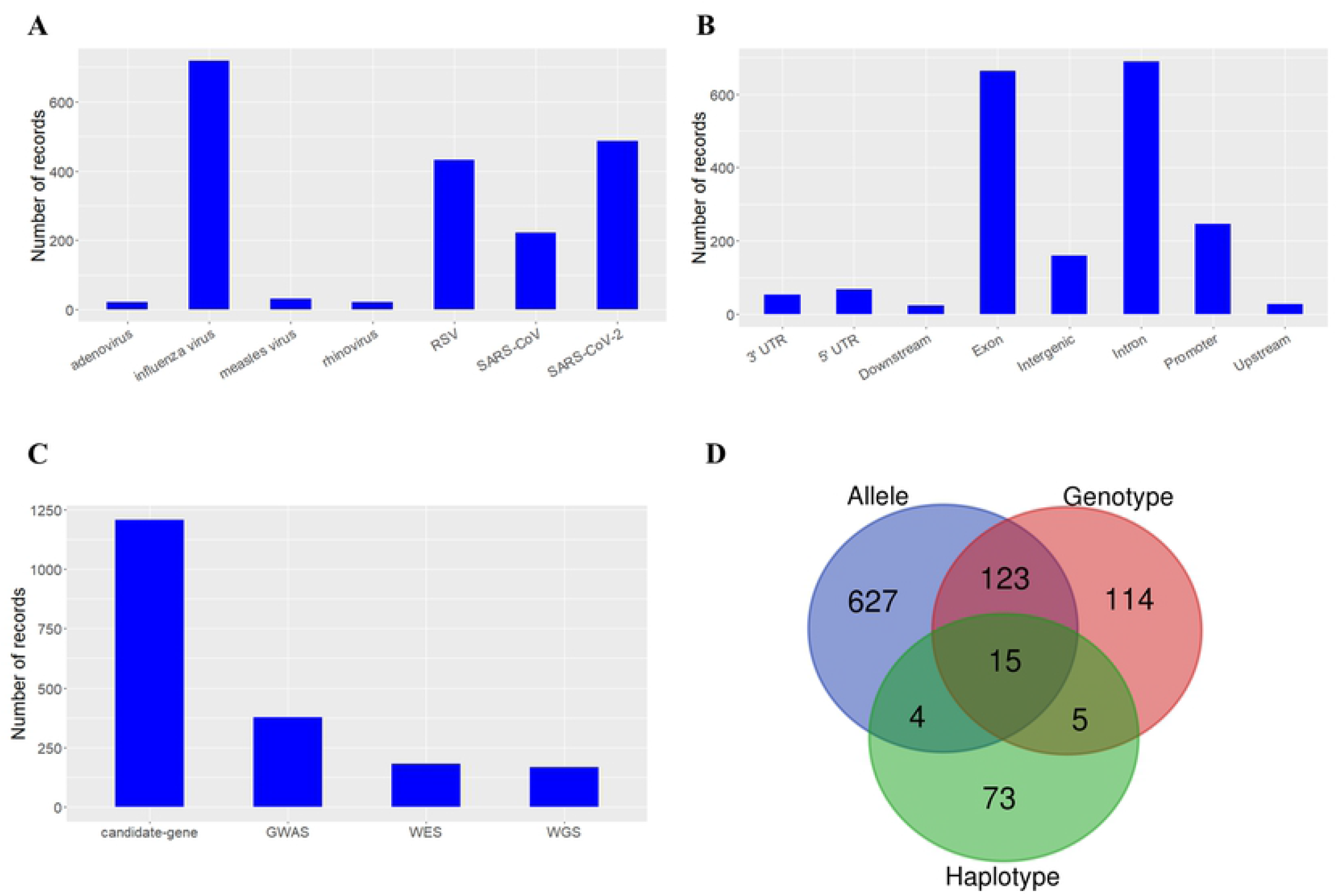

The Network page in the database provides a visual overview of the topological relationship between respiratory viruses and associated genes as shown in Fig 2. Influenza virus, RSV and SARS-CoV-2 have the most associated genes, which is in accordance with these three respiratory viruses having the most study records. On the other hand, a couple of genes are associated with multiple respiratory viruses. Particularly, TNF gene, which is a key mediator of the inflammatory response and is critical for host defense against a wide variety of pathogenic microbes (23), is associated with the greatest number of respiratory viruses.

### GO and pathway analysis of associated genes

We performed GO and pathway analysis of associated genes using Database for Annotation, Visualization and Integrated Discovery (DAVID). The top 10 significantly enriched GO terms and pathways were shown in Table 1 and 2, respectively.

**Table 1.** The top 10 significant GO terms of gene set analysis using associated genes.

| Category | Term | FDR |
| --- | --- | --- |
| GO_BP | immune response | 6.05E-26 |
| GO_BP | inflammatory response | 6.26E-20 |
| GO_MF | cytokine activity | 4.40E-15 |
| GO_BP | positive regulation of inflammatory response | 1.57E-09 |
| GO_BP | positive regulation of T cell proliferation | 2.75E-09 |
| GO_BP | positive regulation of interferon-gamma production | 3.30E-09 |
| GO_BP | innate immune response | 2.29E-08 |
| GO_BP | regulation of complement activation | 4.79E-08 |
| GO_BP | type I interferon signaling pathway | 6.58E-08 |
| GO_BP | lipopolysaccharide-mediated signaling pathway | 6.58E-08 |

**Table 2.** The top 10 significant pathways of gene set analysis using associated genes.

| Category | Term | FDR |
| --- | --- | --- |
| KEGG_PATHWAY | Cytokine-cytokine receptor interaction | 3.78E-19 |
| KEGG_PATHWAY | Influenza A | 4.21E-16 |
| KEGG_PATHWAY | Inflammatory bowel disease (IBD) | 4.21E-16 |
| KEGG_PATHWAY | Herpes simplex infection | 9.03E-16 |
| KEGG_PATHWAY | Measles | 9.66E-15 |
| KEGG_PATHWAY | Rheumatoid arthritis | 1.05E-13 |
| KEGG_PATHWAY | Jak-STAT signaling pathway | 5.57E-13 |
| KEGG_PATHWAY | Leishmaniasis | 5.57E-13 |
| BIOCARTA | Cytokine Network | 8.17E-12 |
| KEGG_PATHWAY | Toll-like receptor signaling pathway | 2.14E-12 |

GO analysis revealed ‘immune response’ and ‘inflammatory response’ were top enriched terms. In addition, other enriched terms such as ‘positive regulation of T cell proliferation’, ‘positive regulation of interferon-gamma production’, ‘regulation of complement activation’, ‘type I interferon signaling pathway’, ‘cytokine activity’ and ‘positive regulation of inflammatory response’ are also related with immune response and inflammatory response, highlighting the central role of these processes against respiratory virus infection (24). Notably, inflammatory response is double-edged sword in respiratory virus infection (25). On one hand, inflammatory response promotes immune response against infection. On the other hand, ‘cytokine storm’ triggered by inflammatory response may worsen the severity of respiratory virus infection (26). Pathway analysis revealed a significant enrichment for pathways directly related to pathogens and autoimmune diseases such as ‘Influenza A’, ‘Inflammatory bowel disease (IBD)’, ‘Herpes simplex infection’, ‘Measles’, ‘Rheumatoid arthritis’ and ‘Leishmaniasis’. There were three enriched pathways related to cytokines, ‘Cytokine-cytokine receptor interaction’, ‘Jak-STAT signaling pathway’ which is the downstream signaling pathway of cytokine interferon (27) and ‘Cytokine Network’. In addition, ‘Toll-like receptor signaling pathway’, which is essential for viral sensing and triggering downstream immune response (28), was also enriched.

## Discussion

To our knowledge, dbGSRV is the first manually curated database containing comprehensive human genetic association information with respiratory viruses. It is composed of several characteristic features worth noting.

First, we thoroughly searched human genetic susceptibility or severity studies of all kinds of respiratory viruses and included them in the database, though most of respiratory viruses were not studied of human genetic susceptibility or severity. More and more studies on a wider range of respiratory viruses are anticipated in the future, and dbGSRV will be updated regularly according to newly available data.

Second, inconsistent results are frequently found in replication studies of genetic association (29, 30), thus only recording positive association results may result in misleading. To provide a more comprehensive and unbiased scope of genetic association studies, we not only included records of positive association results in the database, but also included records of negative results mentioned in the main text of references. Additionally, as conflicting genetic association results might be affected by factors such as ethnicity, sample size, allele frequency and analysis method (31), we also collected information like number, ethnicity and alternate allele frequencies of cases and controls, study type, statistic method, OR, 95% CI and p value for the allele association if available, in order to facilitate users to assess and compare different study results accurately and comprehensively.

Third, dbGSRV provides a user-friendly interface, which offers multiple means for users to query and browse the data, hyper links to access more information of public databases conveniently as well as network visualization of respiratory viruses and associated genes.

In conclusion, dbGSRV will be a convenient resource for researchers to query and retrieve genetic associations with respiratory viruses, which may inspire future studies and provide new insights in our understanding and treatment of respiratory virus infection.

## Acknowledgements

We thank Mr. Tong Chen, Mr. Moyu Liu and Mr. Pu Xue in EHBIO Gene Technology (Beijing) Co., Ltd for their help on the construction of the database

## References

1. Crowe, J. E. Human Respiratory Viruses. Encyclopedia of Virology. 2008:551–8.

2. Kesson AM. Respiratory virus infections. Paediatr Respir Rev. 2007;8(3):240–8.

3. Lafond KE, Porter RM, Whaley MJ, Suizan Z, Ran Z, Aleem MA, et al. Global burden of influenza-associated lower respiratory tract infections and hospitalizations among adults: A systematic review and meta-analysis. PLoS Med. 2021;18(3):e1003550.

4. Li Y, Johnson EK, Shi T, Campbell H, Chaves SS, Commaille-Chapus C, et al. National burden estimates of hospitalisations for acute lower respiratory infections due to respiratory syncytial virus in young children in 2019 among 58 countries: a modelling study. Lancet Respir Med. 2021;9(2):175–85.

5. Wang X, Li Y, Deloria-Knoll M, Madhi SA, Cohen C, Ali A, et al. Global burden of acute lower respiratory infection associated with human metapneumovirus in children under 5 years in 2018: a systematic review and modelling study. Lancet Glob Health. 2021;9(1):e33–e43.

6. Huang C, Wang Y, Li X, Ren L, Zhao J, Hu Y, et al. Clinical features of patients infected with 2019 novel coronavirus in Wuhan, China. Lancet. 2020;395(10223):497–506.

7. Wang C, Horby PW, Hayden FG, Gao GF. A novel coronavirus outbreak of global health concern. Lancet. 2020;395(10223):470–3.

8. Zhu N, Zhang D, Wang W, Li X, Yang B, Song J, et al. A Novel Coronavirus from Patients with Pneumonia in China, 2019. N Engl J Med. 2020;382(8):727–33.

9. Clohisey S, Baillie JK. Host susceptibility to severe influenza A virus infection. Critical care. 2019;23(1):303.

10. Shi T, Vennard S, Mahdy S, Nair H. Risk factors for RSV associated acute lower respiratory infection poor outcome and mortality in young children: A systematic review and meta-analysis. The Journal of infectious diseases. 2021:10.1093/infdis/jiaa751.

11. Zhou F, Yu T, Du R, Fan G, Liu Y, Liu Z, et al. Clinical course and risk factors for mortality of adult inpatients with COVID-19 in Wuhan, China: a retrospective cohort study. Lancet. 2020;395(10229):1054–62.

12. Chapman SJ, Hill AVS. Human genetic susceptibility to infectious disease. Nat Rev Genet. 2012;13(3):175–88.

13. Kenney AD, Dowdle JA, Bozzacco L, McMichael TM, St Gelais C, Panfil AR, et al. Human Genetic Determinants of Viral Diseases. Annu Rev Genet. 2017;51:241–63.

14. Sørensen TI, Nielsen GG, Andersen PK, Teasdale TW. Genetic and environmental influences on premature death in adult adoptees. N Engl J Med. 1988;318(12):727–32.

15. Albright FS, Orlando P, Pavia AT, Jackson GG, Cannon Albright LA. Evidence for a heritable predisposition to death due to influenza. The Journal of infectious diseases. 2008;197(1):18–24.

16. Thomsen SF, Stensballe LG, Skytthe A, Kyvik KO, Backer V, Bisgaard H. Increased concordance of severe respiratory syncytial virus infection in identical twins. Pediatrics. 2008;121(3):493–6.

17. Kwok AJ, Mentzer A, Knight JC. Host genetics and infectious disease: new tools, insights and translational opportunities. Nat Rev Genet. 2021;22(3):137–53.

18. Tahamtan A, Askari FS, Bont L, Salimi V. Disease severity in respiratory syncytial virus infection: Role of host genetic variation. Reviews in medical virology. 2019;29(2):e2026.

19. Di Maria E, Latini A, Borgiani P, Novelli G. Genetic variants of the human host influencing the coronavirus-associated phenotypes (SARS, MERS and COVID-19): rapid systematic review and field synopsis. Hum Genomics. 2020;14(1):30.

20. Lamborn IT, Su HC. Genetic determinants of host immunity against human rhinovirus infections. Hum Genet. 2020;139(6-7):949–59.

21. Pérez-Rubio G, Ponce-Gallegos MA, Domínguez-Mazzocco BA, Ponce-Gallegos J, García-Ramírez RA, Falfán-Valencia R. Role of the Host Genetic Susceptibility to 2009 Pandemic Influenza A H1N1. Viruses. 2021;13(2).

22. Darbeheshti F, Rezaei N. Genetic predisposition models to COVID-19 infection. Med Hypotheses. 2020;142:109818.

23. Waters JP, Pober JS, Bradley JR. Tumour necrosis factor in infectious disease. J Pathol. 2013;230(2):132–47.

24. Chen X, Liu S, Goraya MU, Maarouf M, Huang S, Chen J-L. Host Immune Response to Influenza A Virus Infection. Front Immunol. 2018;9:320.

25. Tavares LP, Teixeira MM, Garcia CC. The inflammatory response triggered by Influenza virus: a two edged sword. Inflamm Res. 2017;66(4):283–302.

26. Fajgenbaum DC, June CH. Cytokine Storm. N Engl J Med. 2020;383(23):2255–73.

27. Ivashkiv LB, Donlin LT. Regulation of type I interferon responses. Nat Rev Immunol. 2014;14(1):36–49.

28. Arpaia N, Barton GM. Toll-like receptors: key players in antiviral immunity. Curr Opin Virol. 2011;1(6):447–54.

29. Chen T, Xiao M, Yang J, Chen YK, Bai T, Tang XJ, et al. Association between rs12252 and influenza susceptibility and severity: an updated meta-analysis. Epidemiology and infection. 2018:1–9.

30. Prabhu SS, Chakraborty TT, Kumar N, Banerjee I. Association between IFITM3 rs12252 polymorphism and influenza susceptibility and severity: A meta-analysis. Gene. 2018;674:70–9.

31. Mozzi A, Pontremoli C, Sironi M. Genetic susceptibility to infectious diseases: Current status and future perspectives from genome-wide approaches. Infection, genetics and evolution: journal of molecular epidemiology and evolutionary genetics in infectious diseases. 2018;66:286–307.

